# Multi-modal data integration for machine learning applications

**DOI:** 10.1101/2025.10.10.681692

**Authors:** Jacques Serizay, Romain Koszul

## Abstract

The integration of multi-modal genomic data, encompassing sequences, annotations, and coverage tracks, remains a major bottleneck in bioinformatics, both for exploratory data analysis and machine learning applications. Current approaches rely on several specialized tools for different data modalities, leading to inefficient workflows and computational overhead. Here, we present momics, a unified framework to consolidate multi-omics data in a single repository and interrogate it with a high-performance query engine. Compared to existing tools, momics ingests genomic sequences, feature annotations, and unlimited coverage tracks into TileDB-backed repositories, and provides a scalable query engine for concurrent multi-modal queries across millions of genomic loci. Our benchmarks demonstrate up to 20-fold better data compression and up to 100-fold speed improvements over standard tools like pyBigWig, with a sublinear time complexity ideal for large-scale queries. Momics provides a python library optimized for exploratory data analysis and machine learning workflows, natively supporting current state-of-the-art bioinformatic ecosystems and cloud storage systems. We demonstrate momics’ utility through two real-world applications: (1) multi-modal data integration of hundreds of ChIP-seq datasets together with genomic sequence, and (2) multi-modal deep learning for chromatin accessibility prediction. By eliminating the need for multiple data parsing tools and providing a unified interface for all genomic data types, momics represents a paradigm shift in how large-scale multi-omics data can be managed and analyzed.

**Key points:**

- momics is a unified framework to consolidate sequences, annotations, and coverage tracks into a single queryable repository, addressing the critical bottleneck in genomic data analysis where researchers must juggle multiple specialized tools for different data modalities.
- We show that momics can achieve up to 20-fold better data compression and 100-fold speed improvements over standard tools, with sublinear time complexity when querying millions of genomic positions simultaneously.
- We use momics to formally demonstrate that multi-modal deep learning models can outperform single-modality approaches in predicting chromatin accessibility, achieving correlation of 0.84 when training with a combination of genomic sequence and MNase data.
- Our results establish a new paradigm for reproducible multi-omics modeling, where entire multi-omics analysis workflows from data storage to machine learning model training can be replicated.

## Introduction

Modern multi-omics profiling technologies generate diverse data modalities including genomic sequences, annotations, and coverage tracks (Baião et al., 2025). These different modalities are often consolidated into an integrated view of the chromatin landscape to understand complex biological systems. However, the current computational infrastructure for multi-omics analysis remains fundamentally fragmented (Luo et al., 2024), and researchers still need to rely on multiple specialized tools and file formats; pyfaidx (Shirley et al., 2015), seqtk (Li, n.d.) or seqkit (Shen et al., 2016) are used to parse FASTA files; bedtools (Quinlan & Hall, 2010) is the swiss-army knife to manipulate BED/GTF files; and pyBigWig (*pyBigWig: A Python Extension for Quick Access to bigWig and bigBed Files*, n.d.) and UCSC utilities (Kent et al., 2010) are the reference to read and write coverage tracks. This proliferation of individual tools, each one dedicated to a certain data modality, leads to important limitations: each query of a multi-modal dataset requires parsing different file formats; data cannot be accessed concurrently across modalities; and software dependencies become increasingly complex as datasets scale to hundreds of tracks.

While the broader data science community has converged on unified frameworks for tabular data (McKinney, 2010; Wickham et al., 2019), and despite efforts to translate this framework for biological data (Hutchison et al., 2024; Open2C et al., 2024), multi-omics studies remain largely constrained by file formats designed for single-modality access. This growing gap between the cutting-edge innovations in data science and the more traditional bioinformatics software stack is increasingly impairing machine learning applications in genomics. Collections of coverage tracks, the main representation of quantitative data generated by international epigenomics consortia such as ENCODE (Sloan et al., 2016), provide an obvious illustration of this issue: large collections of BigWig files can rapidly exceed available virtual memory; pyBigWig specialized parser only supports sequential access patterns to a single track; and the parser cannot execute concurrent queries of multiple tracks. These limitations become prohibitive when training neural networks on multi-modal genomic data, where models require synchronized access to sequences, annotations, and dozens of coverage tracks across millions of genomic loci. Emerging approaches, e.g. keras_dna (*Genomelake: Simple and Efficient Access to Genomic Data for Deep Learning Models*, n.d.; Routhier et al., 2021), now offer the possibility to efficiently stream omics data from different file formats. However, these solutions are specifically tailored to deep learning applications and do not tackle the burden of file management.

Here, we present momics, a unified framework that fundamentally rethinks multi-omics and multi-modal data integration. momics leverages TileDB’s multi-dimensional array infrastructure (Papadopoulos et al., 2016) to consolidate genomic sequences, annotations, and coverage tracks into a single queryable repository. Unlike existing approaches which require separate tools for each data type, momics python API provides a unique high-performance query engine to seamlessly retrieve multi-modal data. This unified architecture enables concurrent queries across hundreds of coverage tracks and millions of genomic positions with several-fold speed improvements over traditional tools, while providing native cloud storage integration for distributed computing environments. We demonstrate how momics transforms multi-omics analysis workflows and accelerates machine learning applications in genomics.

### Momics is a multi-modal data consolidation engine

The momics framework streamlines management of genomic data by ingesting it into a centralized repository, thereby consolidating traditionally disparate file formats into a single, queryable system (**Fig. 1A**). At its core, a momics repository consists of structured hierarchical collections of TileDB arrays (Papadopoulos et al., 2016) organized into three primary data categories: genomic sequences, feature annotations, and coverage tracks (**Fig. 1B**). Upon initialization, momics creates and populates a genome array to store reference sequences, with each chromosome represented as a uni-dimensional TileDB subarray with four attributes (A, T, G, C), indexing genomic positions and their corresponding nucleotides.

**Figure 1.**
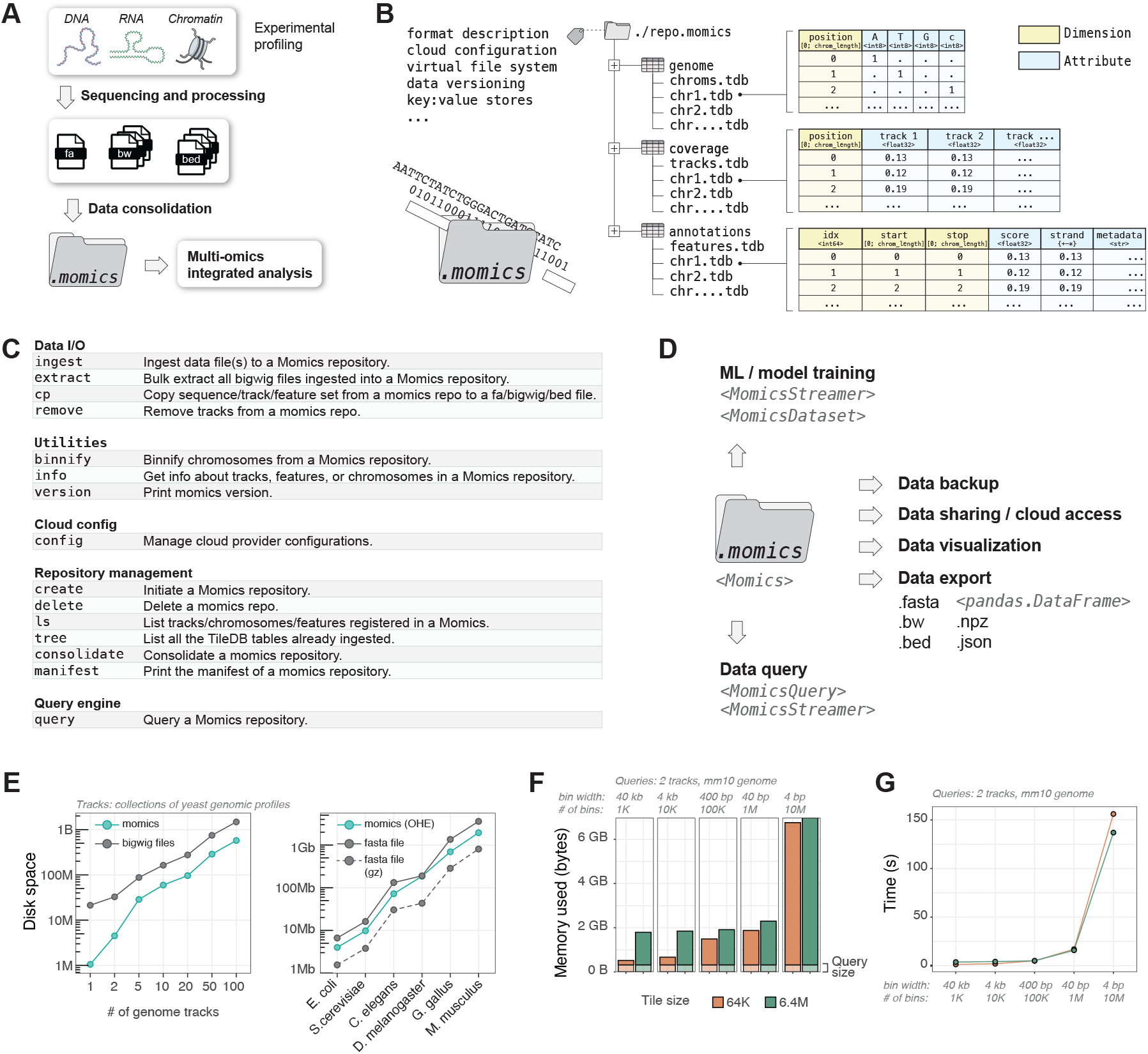
momics framework for multi-modal data integration. ***A***. Schematic overview of the momics workflow. Processed data (DNA, RNA, chromatin) are consolidated into a unified momics repository to enable integrated multi-omics analysis. **B**. Hierarchical organization of a momics repository based on TileDB arrays. **C**. Command-line interface (CLI) utilities for momics repository management. **D**. Classes for efficient data streaming and machine learning application, and utilies for data backup, sharing, visualization, and export in standard formats. **E**. Storage efficiency comparison between momics and traditional bigwig and fasta file formats. **F, G**. Memory (**F**) and time (**G**) usage comparison for different array tile sizes.

The annotations array houses genomic features using three-dimensional subarrays that capture feature IDs, start positions, and end positions, along with optional attributes for scores, strand information, and metadata. Finally, the coverage array contains multi-attribute subarrays, each storing an unlimited number of coverage tracks as column attributes. Metadata can be added to any of the TileDB arrays or subarrays. A command-line interface (CLI) is provided to facilitate the management of momics repositories from a terminal (**Fig. 1C**). Thus, the underlying TileDB hierarchical infrastructure ensures efficient storage of hundreds of quantitative genomic datasets within a single logical structure, transforming the fragmented landscape of genomic file formats into a cohesive data infrastructure where all modalities share a common storage backend.

The true power of momics emerges through its unified query engine, which abstracts away the complexity of accessing different data modalities (e.g. sequence, annotations, and coverage tracks) through a consistent API (**Fig. 1D**). Users can specify genomic regions of interest and retrieve synchronized data across all modalities in a single operation. This is accomplished through the MomicsQuery class, which orchestrates parallel data retrieval from the underlying TileDB arrays and returns results in data science structures like NumPy arrays or Pandas DataFrames (McKinney, 2010). The query engine automatically handles the intricacies of multi-dimensional array slicing and data type conversions that would typically require separate tools and custom scripts for each file format.

The ingestion process leverages TileDB’s configurable compression algorithms, achieving 2- to 10-fold better compression than traditional bigwig or fasta formats (**Fig. 1E**). TileDB arrays are populated using a specific tile size for data storage and compression, with the tile size being the atomic unit retrieved when performing a query. Thus, time and memory required for different query types can be optimized by choosing an appropriate tile size when initially ingesting data into a momics repository. For example, larger tile sizes provide performance improvements for dense query patterns such as millions of small genomic windows (**Fig. 1F**).

Finally, momics’ architecture inherits TileDB’s cloud-native capabilities, enabling repositories to reside on Amazon S3, Google Cloud Storage, or Azure Blob Storage with the same query performance as local storage. This cloud integration is transparent to users, as the same query is used to interrogate a local repository or cloud-hosted public data, and only requires a token-based authentication to query cloud-hosted private repositories. This design enables truly distributed genomic analysis workflows where multiple researchers can concurrently query the same centralized repository without data duplication or synchronization issues.

### Momics is an efficient multi-modal data query engine

momics integrates built-in spatial indexing of multi-modal data stored as TileDB arrays, ensuring rapid random access query. Query performance demonstrates sublinear time complexity, with retrieval times increasing more slowly than the number of genomic intervals queried, maintaining sub-second performance for up to 10 thousand intervals across multiple tracks, corresponding to over 20 million genomic positions (**Fig. 2A**). The system also scales sublinearly when querying increasing numbers of data tracks simultaneously, with only modest time increases when expanding from 2 to 8 concurrent tracks, even at the scale of tens of millions of genomic intervals (**Fig. 2A**). When compared to pyBigWig (*pyBigWig: A Python Extension for Quick Access to bigWig and bigBed Files*, n.d.), the current standard for querying coverage tracks, momics demonstrates superior query performance (**Fig. 2B**). pyBigWig can efficiently handle sequential access to track coverage for a limited number of genomic intervals, but its performance degrades linearly as query complexity increases. In contrast, momics particularly excels in concurrent queries of intermediate numbers of genomic ranges (10K-1M ranges), performing 4-fold (single-threaded mode) and up to 100-fold (multi-threaded mode) faster than pyBigWig implementation, and the performance advantage over pyBigWig grows at increasing scale. When compared to pyfaidx (Shirley et al., 2015), the standard tool for querying genomic sequence, momics achieves 5 to 10-fold speed increases when performing multi-range extraction of tens or hundreds of thousands of one-hot-encoded sequences (**Fig. 2C**). For queries that may exceed memory capacity, momics supports batch processing of large query sets through the MomicsStreamer class, allowing users to specify a batch size to limit memory usage while maintaining efficient retrieval times. This is particularly useful for large-scale machine learning applications, where models may require access to extensive genomic data that cannot be fully loaded into memory.

**Figure 2.**
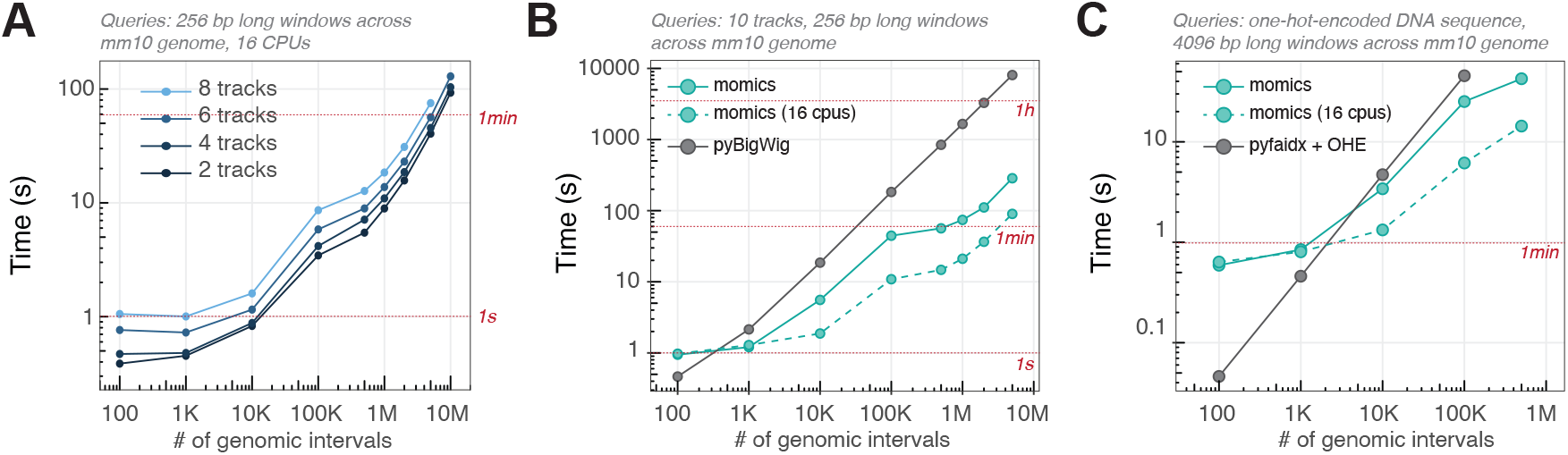
Performance benchmarking of momics against standard genomic tools. **A**. Query time for retrieving coverage data for increasing numbers of 256 bp genomic windows, from 2, 4, 6, or 8 mouse ChIP-seq coverage tracks simultaneously. Red lines indicate 1 second and 1 minute thresholds. **B**. Comparison of momics (single-threaded mode, solid lines; and 16-CPU multi-threaded mode, dashed line) versus pyBigWig for coverage track queries. **C**. Comparison of momics (single-threaded mode, solid lines; and 16-CPU multi-threaded mode, dashed line) versus pyfaidx for genomic sequence extraction. Query times for retrieving one-hot encoded (OHE) DNA sequences from 4096 bp windows across mouse chromosome 10.

Thus, momics ability to handle diverse data modalities, from coverage tracks to sequences and annotations, addresses the growing need for integrated multi-omics analysis frameworks capable of scaling with the exponential growth of genomic data. In addition, data queried from a momics repository can be returned in standard formats like BigWig and fasta, thereby reducing the friction with existing bioinformatic packages and enabling seamless workflow integration. This interoperability, together with superior performance and native cloud support, makes of momics an essential infrastructure for modern genomic data analysis, where traditional file-based approaches fall short.

### Momics facilitates multi-modal data integration for traditional machine learning applications

To illustrate momics’ usefulness for multi-modal data integration in traditional machine learning approaches, we focused on exploring the yeast chromatin landscape by integrating hundreds of ChIP-exo datasets alongside genomic sequences. Using momics, we consolidated the yeast genomic reference sequence and 602 publicly available ChIP-exo datasets (Rossi et al., 2021) (covering various histone modifications and transcription factors) into a single repository. We then interrogated the ingested Sua7/TFIIB ChIP-exo profile to recover width-variable genomic loci enriched for RNA polymerase II (RNAPII) pre-initiation complex (PIC). We next classified these regions using annotations of genes encoding ribosomal proteins (RP), genes bound by SAGA, TUP and/or Mediator/SWI–SNF (STM), genes bound by sequence-specific TFs Abf1 and/or Reb1 (TFO), and unbound genes (UNB). We also annotated loci overlapping with tRNA genes (tRNA) and autonomously replicating sequence consensus sequence (ACS) (Rossi et al., 2021). Finally, we queried the momics repository to extract the average signal of each of the 602 ChIP-exo tracks and the GC and k-mer content (with k from 3 to 7) at these loci.

At this stage, we used different machine learning approaches to characterize genomic loci based on their ChIP-exo profiles and sequence content. First, we performed a PCA on the ChIP-exo profiles and clustered them using a Shared Nearest Neighbor (SNN) approach (**Fig. 3A**). This revealed a clear separation between factors targeting RP or tRNA genes and other gene classes, indicating distinct chromatin signatures associated with ribosomal protein genes. Next, we plotted ChIP-exo scores as a heatmap that visualizes the chromatin landscape across different gene classes (**Fig. 3B**). This further highlighted the unique chromatin features of RP and tRNA genes, dominated by binding of specific transcription factors and chromatin remodelers. Finally, we computed the cumulative distribution function (CDF) of GC content for each gene class (**Fig. 3C**). This highlighted the distinct sequence composition of ACS loci, enriched in AT content. In addition, it revealed a significant difference in GC content between STM genes (median of 37.3%) and TFO or UNB genes (median of 35%), in agreement with previous reports (Rossi et al., 2021).

**Figure 3.**
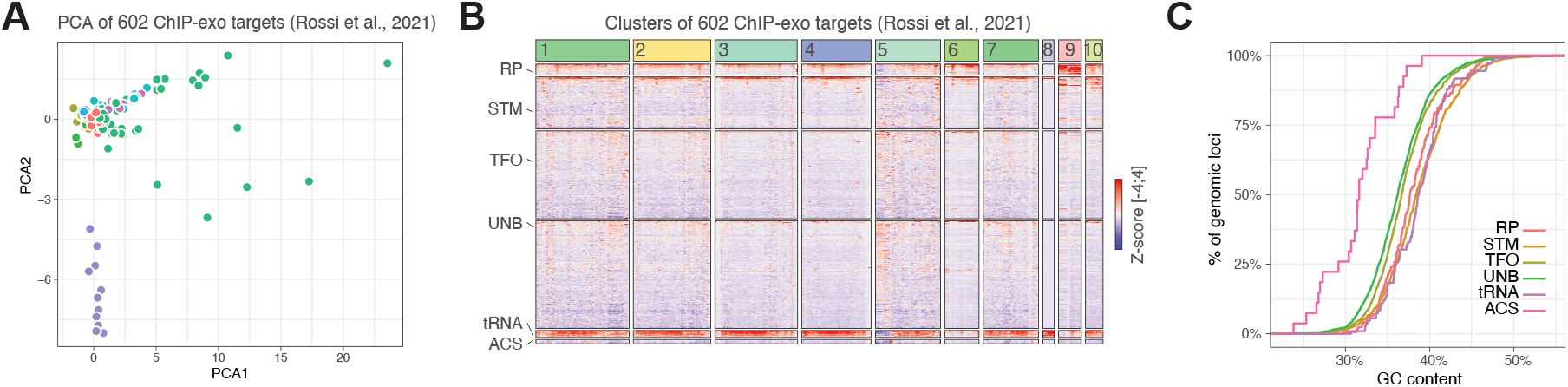
Multi-modal analysis of yeast chromatin landscape using traditional machine learning. **A**. Principal component analysis (PCA) of 602 ChIP-exo targets from Rossi et al. (2021). Points are colored by functional gene categories, revealing distinct clustering of ribosomal protein (RP) and tRNA genes (green) separate from other gene classes along PC1. **B**. Heatmap visualization of ChIP-exo enrichment patterns across the 602 clustered ChIP-exo targets. Rows represent different categories of SUA7+ genomic loci (RP: ribosomal protein genes, STM: SAGA/TUP/Mediator-bound genes, TFO: genes bound by Abf1/Reb1, UNB: unbound genes, tRNA: tRNA genes, ACS: autonomously replicating sequences). Columns show individual ChIP-exo datasets grouped into 10 clusters, with Z-scores indicating enrichment levels. **C**. Cumulative distribution functions of GC content for each gene category.

In conclusion, the ability to seamlessly query and integrate diverse data modalities using momics accelerates the application of traditional machine learning techniques to uncover complex biological insights from multi-omics datasets.

### Momics facilitates multi-modal data integration for deep learning applications

momics extends beyond data management to directly support advanced deep learning applications with the MomicsDataset class, a subclass of Tensorflow Datasets (*TensorFlow Datasets*, n.d.). We demonstrate efficient momics-based NN training by training the multi-modal ChromNN network provided with momics. The ChromNN architecture supports independent input modalities processed in parallel branches, each consisting of concatenated (dilated) convolutional blocks capturing sequential patterns at multiple scales within genomic data (**Fig. 4A**). These parallel input streams undergo independent feature extraction and are eventually concatenated to create a unified latent representation of the multi-omics data. This integrated representation feeds into one or multiple output heads, each predicting a different modality, thereby enabling multi-task learning within a single architectural framework. This design philosophy facilitates the seamless integration of heterogeneous genomic data types for multi-modal predictions, a critical step to investigate complex biological phenomena. MomicsDataset instances can perform batch queries, custom preprocessing, data shuffling and prefetching. Of note, the MomicsDataset class efficiently handles data streaming so that data flows without significant overhead. This ensures that the GPU remains fully utilized during training, even when querying millions of genomic loci across multiple data modalities (**Fig. 4B**).

**Figure 4.**
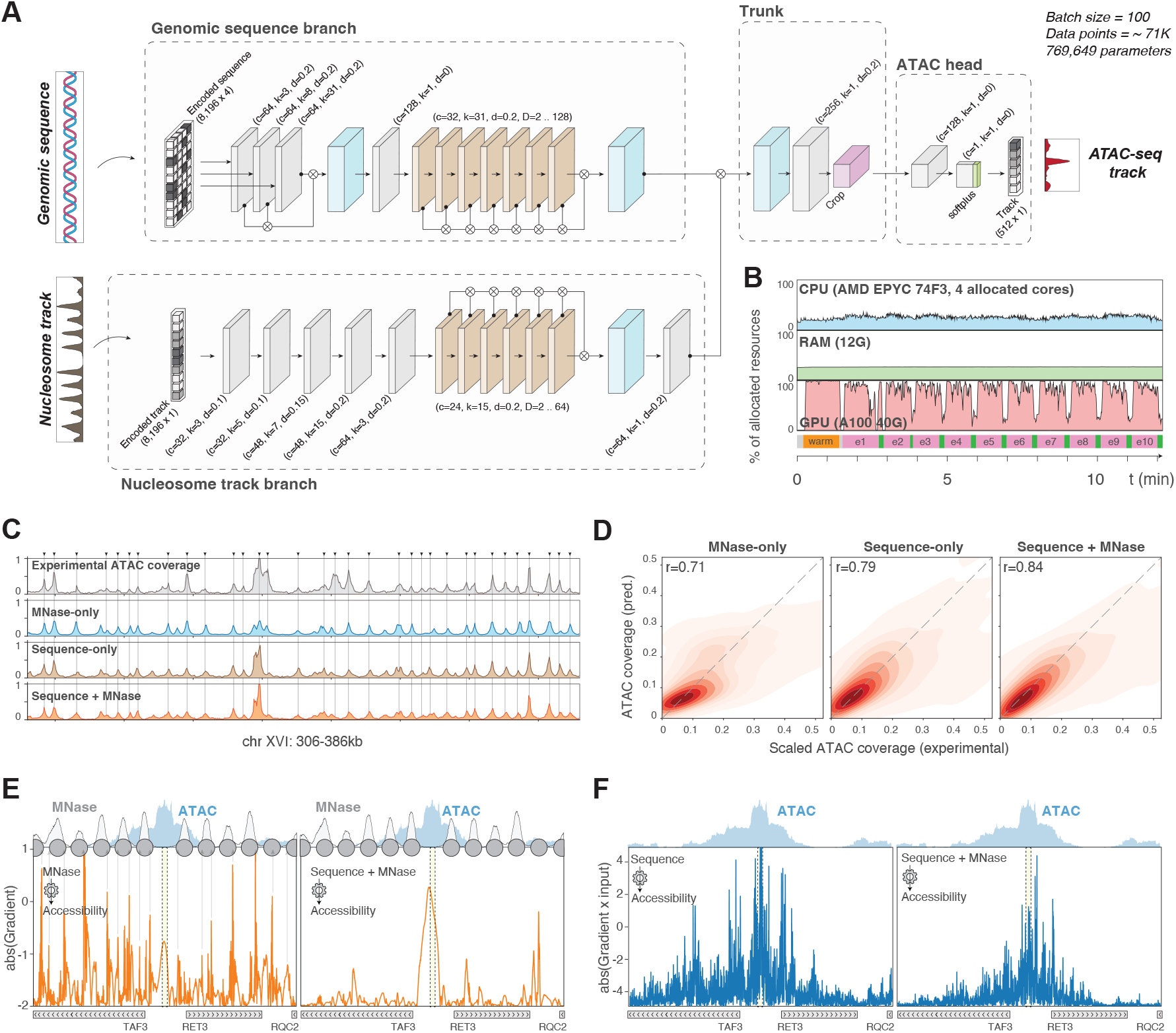
Multi-modal deep learning for chromatin accessibility prediction using momics. **A**.ChromNN neural network architecture for multi-modal chromatin prediction. In this eexample, the model processes two parallel input branches: genomic sequence (one-hot encoded, 8196 bp) and nucleosome track (MNase-seq coverage, 8196 bp). Each branch undergoes independent feature extraction through convolutional blocks before concatenation in a shared trunk network, outputting predicted ATAC-seq signal over 512 bp central regions. **B**.GPU usage during model training demonstrating efficient data pipeline performance. **C**. Predictions from different models on test set, compared with an experimental ATAC-seq coverage (top): MNase-only (blue), sequence-only (green), and combined sequence+MNase model (orange). **D**. 2D density plots showing correlation between experimental and predicted ATAC-seq coverage for the three model variants. Only the test set has been used for these analyses. **E**. Gradient-based attribution analysis for MNase input contribution at the TAF3/RET3 bidirectional promoter. Left: MNase-only model shows high attribution scores at nucleosome boundaries. Right: Combined model focuses attribution within the nucleosome-depleted region (NDR). **F**. Gradient-based attribution analysis for DNA sequence input contribution. Left: Sequence-only model shows distributed attribution across the 2kb region. Right: Combined model restricts sequence contribution to the NDR, demonstrating that multi-modal integration enables the model to focus on the most relevant regulatory features.

To demonstrate *momics’* practical utility for multi-modal predictions, we trained three convolutional neural networks in yeast, to predict chromatin accessibility from different input modalities (**Fig. 4C**). The first model predicts ATAC-seq signal solely from nucleosome scores measured by MNase-seq: 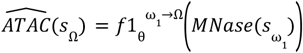, where, s_ω_ denotes a genomic locus of width ω and 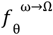 is a neural network that predicts the chromatin accessibility 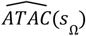 over a locus of width Ωcentered at s, based on the nucleosome scores measured by MNase-seq MNase(s_ω_). The second model uses only the underlying DNA sequence: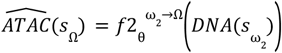 . The third model combines both modalities: 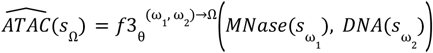. In this example, we set the input length ω to 8,196 and the output length Ωto 512. Compared to previous neural network architectures successfully used to predict individual epigenomic modalities (e.g. nucleosome positioning or RNA Polymerase II binding) from genomic sequence (Meneu et al., 2025; Routhier et al., 2020), ChromNN implements (1) convolutional layers explicitly dedicated to identifying sequence features corresponding to TF binding motifs (8-15 base pairs) and (2) supports multi-modal inputs.

This experimental design allowed us to systematically evaluate the contribution of each data modality to prediction accuracy. We observed different chromatin accessibility predictability for each input modality (**Fig. 4C, 4D**). The model solely trained with MNase-seq data accurately predicted ATAC peak positions, but failed to capture their dynamic range, suggesting that nucleosome positioning alone is not sufficient to dictate the extent of chromatin accessibility. In contrast, the DNA sequence-based model faithfully predicted both the position and height of ATAC peaks, indicating that sequence features encode substantial information about accessibility patterns. Finally, we found that the multi-modal model combining DNA sequence and MNase data outperformed each single-modality model, with a correlation between experimental and predicted profiles of 0.84 compared to 0.71 and 0.79, suggesting that positioning of flanking nucleosomes modulates the accessibility of nucleosome-depleted regions (NDRs) in ways not captured by genomic sequence alone.

To investigate the mechanistic basis of these observations in more detail, we quantified the individual contribution of either the MNase coverage or the genomic sequence for the prediction of chromatin accessibility in regulatory elements. We computed the nucleotide-resolution gradient for each input type (MNase and DNA sequence), in single-modality models or the combined model. This gradient indicates how small changes in the input feature (nucleotide or MNase coverage at each position) would affect the model’s prediction. Over a 2kb locus centered at the TAF3/RET3 bi-dirctional promoter, the MNase gradient in the MNase→ATAC model is characterized by strong absolute scores, reproducibly peaking at nucleosomes borders (**Fig. 4E**, left panel). In contrast, in the combined MNase+sequence→ATAC model, the MNase gradient is peaking at the nucleosome-depleted region (**Fig. 4E**, right panel). This difference suggests whereas the uni-modal model relies on nucleosome positioning to predict accessibility, combining MNase with sequence information allows a bi-modal model to focus on the NDR itself. The same is true for the sequence-based model, which also reveals high absolute scores spread along the 2kb region, whereas the bi-modal model restrains the contribution of the sequence for ATAC prediction to the NDR itself (**Fig. 4F**). This reveals that, while both MNase and genomic sequence both convey enough information to predict chromatin accessibility, the bi-modal model leverages complementary information from both modalities to focus on the most relevant features within regulatory elements, thereby improving the overall prediction of chromatin accessibility.

## Discussion

Momics represent a fundamental shift in how multi-omics data can be stored, accessed, and analyzed. It consolidates diverse genomic data types into unified repositories built on TileDB’s infrastructure, unlocking fast random access and concurrent querying operations. In particular, its flexible, multi-range querying capabilities offer order-of-magnitude performance improvements when compared to traditional file-based approaches, enabling analyses that were previously computationally prohibitive and intractable. This transforms multi-omics data integration from a computational bottleneck into a seamless component of modern genomic workflows.

ChromNN, the multi-modal model we provide with the momics software, builds upon and complements a growing body of research in multi-task chromatin prediction. Models like Borzoi (Linder et al., 2025), Enformer (Avsec et al., 2021), and Sei (Chen et al., 2022) have demonstrated remarkable success in predicting hundreds to thousands of uni-modal chromatin features (e.g. ChIP-seq profiles) from DNA sequence alone. More recent approaches like scooby (Hingerl et al., 2024) have begun exploring true multi-modal integration, for example by predicting combined RNA and ATAC profiles from DNA sequences. However, these modeling achievements often obscure a fundamental challenge: the computational infrastructure required to efficiently manage, query, and integrate the massive multi-modal datasets needed for training. This is where momics makes its critical contribution. While existing studies focus primarily on model architectures and predictive performance, we address the upstream bottleneck of data management that limits the scale and reproducibility of such analyses. Regardless of their internal architecture, momics framework facilitates the training of neural networks by simultaneously querying multi-omics data from a single repository. Furthermore, our work illustrates true multi-modal input integration with ChromNN, where both DNA sequence and MNase-seq data jointly inform predictions, revealing how different data modalities capture complementary aspects of chromatin regulation. By providing the infrastructure to make large-scale multi-modal querying a routine, momics complements these modeling advances and paves the way for even more ambitious integrative studies.

Beyond technical improvements, momics also establishes a new paradigm for reproducible computational biology (Ziemann et al., 2023). Traditional genomics workflows fragment data across multiple file formats and tools, making it difficult to share complete analytical environments. Momics accessible data repositories ensure that complex machine learning analyses can be reproduced, validated, and extended by the broader scientific community. In addition, entire multi-omics datasets can be stored in cloud-hosted repositories and accessed identically by collaborators worldwide.

At the time of writing, the momics framework still faces technical constraints. The TileDB architecture currently requires that multiple coverage tracks be written to chromosomal arrays in a single transaction. To support the incremental ingestion of any number of tracks, momics currently bypasses this limitation by reading back coverage scores already ingested, appending new data, and effectively re-writing the whole array. This linear memory scaling can become limiting for large genomes or extremely large track collections. For example, while 48 Gb of memory is sufficient to ingest 1,000 tracks over the 16 yeast chromosomes in parallel, it can only ingest 100 coverage tracks over a single 120 Mb mouse chromosome.

However, this limitation is actively being addressed by TileDB developers, with the ability to append new attributes without re-writing existing ones expected in the near future.

Looking forward, momics establishes the foundation for a new generation of genomic analyses where data scale is no longer a limiting factor. The framework’s extensibility naturally support expansion to additional data types, and future work on momics could focus on including chromatin conformation capture data, which would complete the multi-omics chromatin landscape. As genomic datasets continue their exponential growth and machine learning becomes central to biological discovery, momics provides a much-needed infrastructure to ensure that computational capabilities keep pace with data generation, ultimately accelerating our understanding of genome function and regulation.

## Material and methods

### Software implementation

Momics is distributed as an open-source Python package under the CC BY-NC 4.0 license. The momics framework is implemented in Python and leverages TileDB arrays, as well as other data science libraries including NumPy, Pandas, and TensorFlow. Momics also supports genomic dataframes through PyRanges (Stovner & Sætrom, 2020). The core functionality leverages the TileDB-Py package to create and manage the underlying TileDB arrays. The MomicsDataset class extends TensorFlow Datasets to facilitate seamless integration with deep learning workflows. The command-line interface (CLI) is built using the Click library, providing an intuitive interface for users to interact with momics repositories. This study was done using momics version 0.4.0, with Python 3.12.0, TileDB-Py version 0.33.0, TileDB core version 2.27.0, NumPy version 1.26.4, Pandas version 2.2.3, TensorFlow version 2.17.1.

### Data sources

The *Saccharomyces cerevisiae* reference genome (R64-1-1) was obtained from the Saccharomyces Genome Database release 104. Genomic annotations were obtained from Supp. Data 1 from Rossi et al., Nature 2021 (Rossi et al., 2021). Blacklist regions correspond to 100bp-long genomic bins overlapping unmappable regions from S. cerevisiae reference genome using 50bp reads and default bowtie2 (Langmead & Salzberg, 2012) mapping. Stranded ChIP-exo tracks were downloaded from https://www.datacommons.psu.edu/download/eberly/pughlab/ and merged into unstranded ChIP-exo tracks. Chromosome names were converted to Ensembl format to match the reference genome, and replicates were averaged when available using pyBigWig. Tracks were then normalized by clamping values between the 0 and 99.9 percentiles, followed by min-max rescaling to [0, 1]. ATAC-seq, Scc1 ChIP-seq, MNase-seq and RNA-seq profiles were retrieved from Meneu, Chapard, Serizay et al., Science 2025 (Meneu et al., 2025) (ATAC: GSM8640705; MNase-seq: GSM8640795; RNA-seq: GSM6703673; Scc1 ChIP-seq: IP GSM6703640, Input GSM6703641).

### Performance benchmarking

To compare the disk space used between bigwig tracks and momics repositories, we used 1–100 ChIP-exo tracks (see above). To compare the disk space used between fasta files and momics repositories, we compared the size of the fasta file (gzipped or unzipped) was compared to the size of the momics repository after ingestion of the genome reference sequence file. Genome reference sequence for *E. coli* was obtained from Ensembl release 62. Genome reference sequences for *C. elegans, G. gallus, D. melanogaster*, and *M. musculus* were obtained from Ensembl release 115. To check the memory used by different queries, we created two momics repositories with different tile sizes (64KB and 6.4MB) and ingested two ChIP-seq tracks obtained from ENCODE data portal. We then performed queries for a fixed total number of 80M (40M from each track) genomic positions, varying the number of non-overlapping genomic bins and bin widths accordingly. We monitored RAM usage with memory-profiler. To check the time spent by different queries, we created a momics repository with a 64KB tile sizes (default) and ingested ten mouse ChIP-seq tracks obtained from the ENCODE data portal (ENCFF707HHX, ENCFF163SBS, ENCFF880KKC, ENCFF763GCB, ENCFF857GJE, ENCFF097KTK, ENCFF531JKE, ENCFF705OWT, ENCFF643WMY, and ENCFF550KLM). We then performed multi-threaded queries (16 CPU) for a varying number of non-overlapping 256bp-long genomic windows (100 to 10M) and for a varying number of ChIP-seq coverage tracks (2, 4, 6, 8). To compare the time spent for different coverage queries between pyBigWig and momics, we used the same momics repository and queries, but retrieved data for the full set of ten tracks. To compare the time spent for different sequence queries between pyfaidx and momics, we used the a momics repository after ingesting *mm10* genome reference. We performed single- or multi-threaded queries (16 CPU) for a varying number of non-overlapping 4096bp-long genomic windows (100 to 500K). When querying sequences with pyfaidx, we included the time needed to perform on-the-fly one-hot-encoding.

### Traditional machine learning analysis

To demonstrate momics usability for traditional machine learning analysis, we ingested the 602 ChIP-exo tracks into a momics repository (see above), together with the yeast genomic sequence. We identified SUA7-positive loci as genomic windows longer than 100-bp with an average score > 0.1, non-overlapping with blacklist regions (see above). We then used momics to extract the one-hot-encoded sequence and per-nucleotide ChIP-exo coverage for each of the 602 track over the SUA7+ loci. We computed GC and kmer occurrences using numpy. We exported these results as csv files and performed machine learning applications after importing them in R, using tidyverse and tidyomics. We annotated the SUA7+ loci based on genomic annotations retrieved from Supp. Data 1 from Rossi et al., Nature 2021 (Rossi et al., 2021). We performed PCA using BiocSingular and SNN clustering using igraph with k=6 (Csárdi & Nepusz, 2006). We visualized ChIP-exo scores per genomic loci as a heatmap using ComplexHeatmap (Gu et al., 2016). We visualized PCA and CDF curves using ggplot2 (Wickham et al., n.d.).

### Deep learning for chromatin accessibility prediction

We trained three models to predict ATAC coverage in yeast based on ChromNN architecture provided in momics: one with a single “nucleotide” input modality, one with a single “MNASE” input modality, and the last with both modalities. We used a feature size of 8192, a target size of 512, and a batch size of 500. We used tiling genomic bins obtained from chromosomes 1 to 13 (stride of 48) for training, from chromosomes 14 and 15 for validation, and from chromosome 16 for testing. ChromNN models were compiled using an Adam optimizer, a learning rate starting at 0.001 and gradually decreasing up to 1e-5, a loss function defined as the sum of MAE and 1-Pearson’s correlation. An early stopping procedure was applied during training to prevent models from overfitting. The loss function was calculated on the validation set at every epoch to evaluate the generalisability of the model. The training procedure was stopped if the validation loss did not decrease at all for 6 epochs and the model parameters were set back to their best performing value. The training procedure was performed on an HPC compute node with 4 allocated cores from an AMD EPYC 74F3 CPU, 12 GB allocated RAM, and exclusive access to an NVIDIA A100 40GB GPU, and typically converged within 15-20 epochs and under 2h. After training, Pearson’s correlation was calculated by predicting ATAC coverage over the test dataset (bins over the chromosome 16).

### Model explainability

To get insight into the decision-making process of the trained models, we computed nucleotide-resolution gradients for each input type (MNase and DNA sequence), in single-modality models or the combined model. For a given input and model, we computed the gradient of the output with respect to the input using TensorFlow’s GradientTape functionality. We then took the absolute value of these gradients to assess the magnitude of influence each input feature has on the output prediction.

## Author contributions

Jacques Serizay (Conceptualization, Methodology, Software, Formal analysis, Investigation, Resources, Writing—Original Draft, Writing—other versions, Visualization) and Romain Koszul (Supervision, Project administration, Funding acquisition).

## Conflict of interest

None declared.

### Acknowledgments

We thank all our colleagues from the laboratory Régulation Spatiale des Génomes for fruitful discussions on the work. Funding: This work was funded, in whole or in part, by grants from the French government, managed by the Agence Nationale de la Recherche under the France 2030 programme to R.K. [ANR-23-CHBS-0002; ANR-22-CE12-0013; ANR-19-CE13-0027-02]. J.S. was supported by an ARC fellowship. A CC-BY public copyright license has been applied by the authors to the present document and on all subsequent versions up to the author-accepted manuscript version arising from this submission, in accordance with the grants’ open-access conditions.

## Data and code availability

The code for momics is publicly available at https://github.com/js2264/momics. momics can be installed from PyPI (https://pypi.org/project/momics/). Documentation and tutorials are available at https://js2264.github.io/momics.

All data presented in this manuscript have already been published. Stranded ChIP-exo tracks were downloaded from https://www.datacommons.psu.edu/download/eberly/pughlab/. Mouse data (ENCFF707HHX, ENCFF163SBS, ENCFF880KKC, ENCFF763GCB, ENCFF857GJE, ENCFF097KTK, ENCFF531JKE, ENCFF705OWT, ENCFF643WMY, and ENCFF550KLM) was retrieved from the ENCODE data portal. ATAC-seq, Scc1 ChIP-seq, MNase-seq and RNA-seq profiles were retrieved from Meneu, Chapard, Serizay et al., Science 2025 (Meneu et al., 2025) (GSM8640705, GSM8640795, GSM6703673, GSM6703640 and GSM6703641).

